# Resolving Competition in Auditory Cortex: Effects of Emotional Content and Misophonia Sensitivity

**DOI:** 10.1101/2025.09.22.677819

**Authors:** Laura Ahumada, Faith E. Gilbert, Richard T. Ward, Jourdan J. Pouliot, Ryan P. Mears, Andreas Keil

**Affiliations:** Laboratory for the Study of Brain, Body, and Behavior, University of Florida; Department of Psychology, University of Florida

**Keywords:** Attentional Competition, Auditory Steady-State Response (ASSR), Emotional Content, Misophonia

## Abstract

How the human auditory cortex prioritizes relevant information amid concurrent sounds has been a long-standing question in auditory cognitive neuroscience. The present study used auditory steady-state responses (ASSR) to tag the electrocortical response to a tone embedded in concurrent naturalistic sounds, addressing methodological challenges with overlapping auditory streams. Participants endorsing low (LMS) or high in misophonia symptoms (HMS)—a condition with decreased tolerance to specific, typically orofacial, sounds—were recruited. Sounds varied in their emotional valence (pleasant, neutral, unpleasant, and orofacial) to investigate how emotional content modulates attentional competition and how competition is resolved in listeners with misophonia traits. Affective ratings, alpha-band changes, and pupil dilation in response to the sounds were also assessed. Hypothetical models of competition were tested, revealing a facilitation trend in the ASSR amplitude when accompanied by pleasant and unpleasant, compared to neutral sounds, regardless of misophonia symptoms. However, ASSR was selectively reduced in the HMS but not the LMS group when accompanied by orofacial sounds. Analyses of alpha-band, pupil, and rating data showed that the groups differed primarily in their response to pleasant sounds and orofacial sounds, with the HMS group exhibiting a stronger response to orofacial sounds than the LMS group.

---

The human brain has evolved mechanisms for managing limited capacity systems by prioritizing stimuli that are relevant for behavior, while ignoring less relevant sensory information (Desimone & Duncan, 1995). These mechanisms, often referred to as selective attention, are most apparent when people confront complex environments with many concurrent streams of information that overlap in space and time (Kastner & Ungerleider, 2001; Mather & Sutherland, 2011). The selection of one of many concurrent stimuli or stimulus features may be guided by properties of the stimulus (such as their emotional salience, Todd & Manaligod, 2018), by experimental instructions (Desimone & Duncan, 1995), or both (Boylan et al., 2019). In the visual system, studies of competition and its resolution by selective attention often capitalize on the principles of visuocortical organization, including its retinotopy (Clark & Hillyard, 1996; Green et al., 2017), or feature specificity (Andersen et al., 2011; McMains et al., 2007).

Examining prioritization of one sound among many concurrent sounds is, however, often made difficult by the fact that their spatial sources may be overlapping. This is especially true in everyday listening situations, and when the cortical representation areas of complex auditory features are not distinct or known a-priori (Leaver & Rauschecker, 2010). This issue also challenges discriminating the neural responses that result from the attentional selectivity to a particular auditory stimulus. Thus, new methodological strategies are necessary to evaluate interactions between concurrent, overlapping streams of auditory information.

Previous research has evaluated auditory competition using a variety of tasks (Hymel et al., 2000; Scanlon et al., 2019; Sussman et al., 2003). For instance, using an oddball paradigm, Sussman and colleagues (2003) found that the mismatch negativity (MMN) response from an unattended auditory channel disappeared when a target detection task was introduced in the opposite auditory channel. In contrast, adding a non-detection task did not produce this effect, suggesting that auditory competition during concurrent binaural streaming arises from top-down processes (Sussman et al., 2003). In addition, the brain’s intrinsic oscillatory activity has been also investigated during competition. Specifically, in the context of speech extraction under noisy environments, changes in attentional demands markedly modulated the alpha frequency band - 8-13 Hz - (Obleser & Weisz, 2012; Strauß et al., 2014; Wöstmann et al., 2017). Under these contexts, gradually increasing the acoustic detail of a speech distractor while attending to a simultaneous speech target increased apha- band power (Wöstmann et al., 2017). This effect has been interpreted as a mechanism of selective inhibition of task irrelevant/noisy information (Strauß et al., 2014; Wöstmann et al., 2017). Therefore, auditory alpha-band enhancement reflects a modulatory response that involves the relative disengagement of cortical networks from the processing of irrelevant or distracting events (Foxe & Snyder, 2011).

Although the experimental designs discussed above are helpful for elucidating selection and prioritization mechanisms, none of them allows researchers to directly and independently quantify the neural response to a specific auditory stream that is embedded in two or more concurrent, overlapping streams. Steady-state potentials are a method for addressing this issue (Wieser et al., 2016).

Specifically, the steady-state potential frequency tagging method capitalizes on the fact that periodic stimulation of a sensory system results in a strong neural response at the same temporal rate (Norcia et al., 2015; Picton et al., 2003). Thus, when presenting multiple stimuli at distinct modulation rates, the resulting periodic response for each stimulus can be measured in the frequency domain, and separated from responses to other stimuli (Matulyte et al., 2024). In the case of steady-state visual evoked potentials (ssVEP), a robust amplitude enhancement has been observed when a tagged stimulus is selectively attended, or when it is motivationally relevant because of its content or predictive value (Andersen et al., 2008; Friedl & Keil, 2021; Moratti et al., 2006; Morgan et al., 1996). Similar findings have been obtained in the auditory system (Bharadwaj et al., 2014; Müller, 2009; Skosnik et al., 2007), where auditory stimuli are typically modulated at fixed temporal rates to evoke auditory steady-state responses (ASSRs).

Compared to ssVEPs, ASSRs tend to be more challenging to measure because the configuration of the primary auditory cortex (A1) and adjacent cortical areas tends to project to relatively distal EEG electrodes (Weisz et al., 2004). Additionally, both ongoing endogenous brain oscillations and different types of noise may overlap with the ASSR signal (Picton et al., 2003). Computational approaches for maximizing the signal-to-noise ratio (SNR) and effectively isolating the steady-state response are therefore particularly useful for studies with ASSRs. The rhythmic entrainment source separation (RESS; see Cohen & Gulbinaite, 2017) is one such approach, which uses linear spatial filtering techniques to maximize an oscillatory signal.

## The present study

In the present study, we use the RESS algorithm to isolate the ASSR evoked by a tone that was simultaneously presented with naturalistic sounds. In order to evaluate how the overlap between two spatially and temporally concurrent auditory streams affects tone processing, we manipulated the content of the naturalistic sounds in terms of their emotional relevance. Changes in the tone-evoked ASSR amplitude were assessed as a function of the accompanying naturalistic sounds. Additionally, to examine inter-individual differences in the motivational relevance of naturalistic sounds, we examined a sample of individuals scoring low and high in misophonia symptoms in the study.

Misophonia is a disorder characterized by a decreased tolerance to certain sounds that are usually generated by another person, such as loud chewing, heavy breathing, and sniffing (Swedo et al., 2021). Individuals with this condition report experiencing high levels of distress when they are exposed to these sounds (Cavanna & Seri, 2015; Jager et al., 2020). Commonly, sufferers report experiencing intense feelings of disgust, frustration, and anger when exposed to trigger sounds (Edelstein et al., 2013) to an extent that impairs functioning (Aazh et al., 2019). Thus, orofacial sounds are expected to possess heightened motivational relevance for listeners that endorse misophonia symptoms and affect competitive interactions more strongly than other categories of sound.

Based on the literature discussed above, we aim to use the frequency specificity of the ASSR to measure how the auditory cortical processing of a tone stimulus is affected by a concurrent naturalistic sound, varying in content. Existing theoretical models of competition between stimuli varying in motivational relevance provide different predictions: Under a <u>biased competition</u> (interference) perspective, the significance of a motivationally relevant stimulus, conveyed through instructions (Carrasco, 2011), or through arousing stimulus content (Mather & Sutherland, 2011) will bias processing towards the relevant stimulus or feature, typically causing an <u>interference</u> of responses to concurrent features of stimuli (De Echegaray et al., 2024; Wieser et al., 2012). In the present paradigm, the interference framework would thus predict that emotionally engaging content (e.g., pleasant and unpleasant sounds) prompts an attenuation of the ASSR amplitude evoked by the concurrent tone (Matulyte et al., 2024).

Alternatively, several theoretical models and empirical findings suggest that the auditory cortex is capable of synergistically amplifying multiple streams (Bidet-Caulet et al., 2007; Riels et al., 2020), or selecting a complex auditory object (Shinn-Cunningham, 2008), which may contain several features or sub-streams (Alain & Arnott, 2000; Lazzouni et al., 2010; Manting et al., 2020). In this viewpoint, capacity limitations appear later in processing (Holmes et al., 2021; Tóth et al., 2019), or are manifest only in intermodal paradigms where multiple auditory and visual streams compete under high perceptual load (Matulyte et al., 2024). This suggests an alternative hypothesis, which is that the presence of motivationally relevant sounds may alert listeners to the auditory modality, or to the compound object consisting of the tone and the sound (Alain & Arnott, 2000). This <u>facilitation</u> model predicts that the tone-evoked ASSR amplitude is heightened when accompanied by an engaging (pleasant, unpleasant) naturalistic sound.

We tested these alternative hypotheses using a naturalistic listening situation in which no cognitive task was employed. To characterize emotional engagement with the sounds and to allow manipulation checks, pupil dilation during listening and ratings of hedonic valence and emotional arousal were measured. Based on the literature mentioned above, changes in the EEG alpha-band were also examined. EEG alpha power decreases over central and parietal cortices during some complex listening tasks, and has been linked to the successful parsing of speech (Obleser & Weisz, 2012) and active listening (Ala et al., 2020; Wisniewski & Zakrzewski, 2023). These measures served as additional variables for supplementing the ASSR measurement, and as complementary evidence for and against the interference versus facilitation hypotheses.

## Methods

### Participants

In total, 64 university students from the University of Florida and community-based participants took part in this study, see Table 1 for demographics information. Participants completed a prescreening using the Misophonia Symptom Scale (MSS) to include subjects who exhibit strong traits of misophonia symptoms (Misophonia Questionnaire – MQ; Wu et al., 2014). Upon arrival, participants read and signed the informed consent form for this study. They indicated having no family history of epilepsy and they confirmed having normal or corrected-to-normal vision. Upon completion of the study, participants were compensated with course credits, in the case of the university students, or a monetary compensation of $20 USD, in the case of the community-based participants. This study was reviewed and approved by the University of Florida Institutional Review Board in accordance with the Declaration of Helsinki.

**Table 1.** Demographic Information.

|  | Total | LMS | HMS |
| --- | --- | --- | --- |
| **Sex** |  |  |  |
| Male | 18 | 8 | 10 |
| Female | 46 | 26 | 20 |
| Total | 64 | 34 | 30 |
| **Gender** |  |  |  |
| Man | 18 | 8 | 10 |
| Woman | 44 | 25 | 19 |
| Non-binary | 2 | 1 | 1 |
| **Race** |  |  |  |
| White | 56 | 29 | 27 |
| Black or African American | 2 | 1 | 1 |
| Asian | 3 | 1 | 2 |
| Other | 2 | 2 | 0 |
| Prefer not to answer | 1 | 1 | 0 |
| **Ethnicity** |  |  |  |
| Not Hispanic | 53 | 27 | 26 |
| Hispanic | 11 | 7 | 4 |
| **Age (mean)** | 20 | 21 | 20 |
| **MSS score (mean [SD])** | 10.83 [6.02] | 6.35 [2.72] | 15.90 [4.50] |

*Note.* Demographic information is reported for the total participant sample, as well as for participants sorted into the LMS group and HMS group. Mean and the standard deviation of the mean (SD) are also reported for the MSS scores when participants scored low in misophonia symptoms (< 12, LMS), and high in misophonia symptoms (>= 12, HMS).

### Auditory Stimuli

Complex tones were implemented in the Matlab environment, version 2017b. The tones consisted of pure 440 Hz sine waves with a duration of 6 seconds, sampled at 22 kHz. The tones were amplitude-modulated (99% depth) by means of a sinusoid function at 41.2 Hz. The modulation frequency was used to elicit the auditory steady-state response (ASSR). In addition, naturalistic auditory stimuli from the International Affective Digitized Sounds System (IADS-2) were utilized (Bradley & Lang, 2007) to be presented simultaneously with the tones. 9 neutral, 9 pleasant, 9 unpleasant sound categories were curated from the IADS-2. Furthermore, 9 human orofacial sounds (frequent misophonia triggers) were created, based on exemplars retrieved from various online sources. The sound files were normalized to the same sound energy (root mean square) as the IADS sounds and cut and sampled to match their length. Sounds were prioritized that had sustained orofacial sounds, rather than transient, events. They included auditory scenes such as messy apple eating, speaking with full mouth, and sustained slurping. A list of the sounds used in the experiment can be found in Table S1 (Supplementary Material section). The resulting 36 naturalistic sounds were presented twice each, in two separate blocks (one without and other with sounds’ ratings after each trial). Together, both blocks had 18 trials per condition, which summed up to a total of 74 trials. The tones and sounds were delivered through two studio monitor speakers (Behringer Studio 50) located behind the participants.

The sound pressure level in the experimental chamber was measured using a sound level meter (RisePro) and maximum sound intensity level, measured 100 cm from the speakers, was set to 70 dB(A). The root mean square (RMS) of all naturalistic sounds was set to be matched across the four content conditions (pleasant, neutral, unpleasant, and orofacial). To accomplish this, sound time series were z-transformed first and multiplied with an amplitude factor until the RMSs did not differ between stimuli.

### Self-Assessment Manikin (SAM)

The Self-Assessment Manikin (SAM; Bradley & Lang, 1994) was employed to assess self-reported emotional arousal and hedonic valence (pleasure/displeasure) for the naturalistic sounds used in the experiment. For the arousal assessment, a scale from 1 to 9 was used, in which 1 was used to indicate that the sound was calm and 9 to indicate that it was highly arousing. A similar scale was utilized for the valence assessment, in which 1 was used to indicate that the sound was highly pleasant and 9 to indicate that it was highly unpleasant.

### Auditory competition paradigm

The experimental paradigm consisted of two blocks of trials. In the first block, a trial started by presentation of a tone by itself for a duration of 1000 ms. Then, the naturalistic sounds was played together with the tone for 5000 ms until the tone and the accompanying sound co-terminated (see Figure 1). Sounds were randomized such that each of the 36 naturalistic sounds were presented once during the block. The inter-trial interval (ITI) was set to 7400 ms. While listening to auditory stimuli, participants were instructed to fixate on a fixation dot on a Display++ LCD Monitor (Cambridge Research Systems Ltd., Rochester, UK), which spanned a 0.18° visual angle. The second block was similar to the first block, except that arousal and valence ratings were invited at the end of each trial.

**Figure 1.**
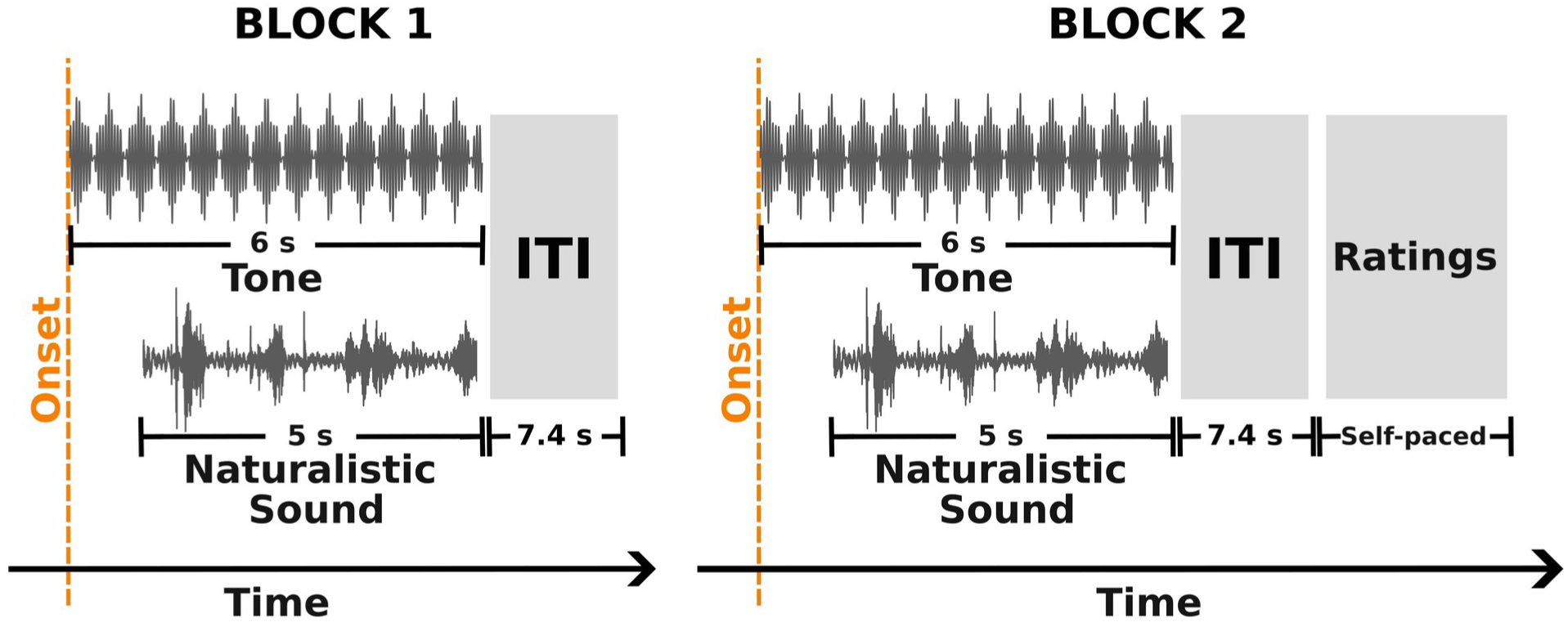
Trial Design per Block. *Note.* The auditory competition paradigm involves presenting a 440 Hz tone (modulated at 41.2 Hz) followed by a naturalistic sound, with a 1-second delay between the two. Both auditory streams terminate simultaneously, 6 seconds after tone onset. The inter-trial interval is set at 7.2 seconds. The experiment consists of two blocks: in the first block, participants are presented with the auditory streams alone, while in the second block, participants rate the arousal and valence of the naturalistic sounds after each trial.

### Loudness discomfort level (LDL) assessment

Misophonia has often been linked to hyperacusis, a condition in which listeners display low tolerance to sounds and strong aversive responses to most auditory stimuli (Jastreboff & Jastreboff, 2023). Thus, we aimed to assess loudness discomfort levels (LDLs) to examine the contribution of hyperacusis to misophonia scores in the present sample. To assess LDLs, participants were presented with a series of one-second tones at five different pitches (320, 544, 925, 1,572, and 2,673 Hz), with a maximum of 10 volume levels for each pitch. After a tone was played, participants were asked if they would be able to tolerate the tone at a slightly higher volume. If they agreed, the tone was played again at a higher volume; if not, a tone at another pitch was played and the ramp-up procedure began at a minimal loudness level for this pitch. The volume levels, measured with an audiometer, ranged from 69 to 91 dB(A) and increased in steps of 2.5 dB. The test continued until participants reached the maximum volume level for each pitch. Hyperacusis sensitivity was measured by summing up the loudness levels across all pitches (from 1 to 50), with higher scores indicating greater auditory tolerance. Although this assessment provides valuable insights into hyperacusis sensitivity, it is essential to note that it does not offer a comprehensive evaluation of the condition. In addition, four participants withdrew during the test.

### Procedure

After reading and signing the consent form, participants completed for a second time the MSS (Wu et al., 2014). The questionnaire scores were used later to separate those individuals who endorsed a large number of misophonia symptoms (high misophonia symptoms or HMS) from those who endorsed minimal symptoms (low misophonia symptoms or LMS). After filling in the questionnaire and completing the hyperacusis test, participants were directed to the experimental chamber. In this chamber, the experimenter explained the nature of the study and reminded participants to maintain fixation in the middle of the screen and avoid excessive body movements. Participants were informed that there may be a variety of sounds ranging in pleasantness but were not informed about the misophonic characteristic of some sounds. Subsequently, the eye tracker was calibrated and validated using a 12-point grid. Then, the sensor net was applied, EEG and pupil recordings started, and the experimental session began. After the experiment, the net was removed and rinsed. Any additional questions participants had about the experiment were answered and they were thanked for their participation.

### EEG pre-processing

A Net Amp 300 amplifier (Magstim EGI, Oregon, US) together with 128-channel HydroCel Geodesic sensor nets were used for EEG recording. Data were recorded at a sample rate of 500 Hz, with reference electrode at Cz. Impedances were kept below 60 KΩ. Later, data were pre-processed using a Matlab (vs. 2022, Natick, MA, USA) pipeline that combines EEG processing functions from EEGLAB and Emegs2.8 (Delorme & Makeig, 2004; Peyk et al., 2011). Butterworth filters, with cut-off points defined as the 3 dB point, were applied offline: High-pass filter at 4 Hz (3^rd^ order) and low-pass filter at 55 Hz (30^th^ order). Eye movement correction was applied using an automated regression-based algorithm, which used the electrical eye activity from the horizontal and vertical EOG electrodes to remove EOG artifacts (Schlögl et al., 2007). Data segmentation was then set for 600 ms before and 6000 ms after tone onset, which resulted in a data structure of electrodes by time and by trials. Artifact-affected trials were identified following the Statistical Control of Artifacts in Dense Array EEG/MEG Studies (SCADS; Junghöfer et al., 2000). Three parameters from the electrodes and trials’ distributions were calculated: The median absolute voltage amplitude, the standard deviation of the voltage, and the largest transient value in a given trial and channel. These parameters were standardized and linearly combined to obtain a quality index per electrode, electrodes by trials, and trials. Electrodes that had a quality index at 2.5 standard deviations above the median of the resulting parameters’ distribution, were interpolated using spherical splines. Similarly, trials which quality index surpassed 1 standard deviations above the median of the parameter’s distribution were rejected. The numbers of trials included per condition after artifact rejection did not differ betwee pleasant (15.8), neutral (15.9), unpleasant (16.0), and orofacial (15.8). Lastly, all voltage data were re-referenced to the average.

### RESS (Rhythmic entrainment source separation) and ASSR (auditory steady-state response)

The rhythmic entrainment source separation (RESS) is a multivariate source separation method that isolates and maximizes the signal-to-noise ratio of a signal of interest (Cohen & Gulbinaite, 2017), in this case the auditory steady-state response. RESS is based on the generalized eigendecomposition, in which two covariance matrices are built to obtain the eigenvector with the largest eigenvalue. One covariance matrix is obtained for the frequency of interest (signal) and the other for the two adjacent frequencies (reference signal). The resulting eigenvector with the largest eigenvalue is used then as the best linear spatial filter to weight the raw data and so maximize the signal of interest (Cohen & Gulbinaite, 2017).

For this study, the signal of interest was the tone modulation frequency at 41.2 Hz, and the lower and upper adjacent frequencies corresponding to 40.2 and 42.2 Hz, respectively. A frequency-domain Gaussian-shaped filter was used to isolate the spectral power of the signal and reference signal’s frequencies. For this purpose, the filterFGx function supplied together with the RESS functions (Cohen & Gulbinaite, 2017) was employed. The time window of analyses for the frequency-domain filter started at 200 ms after tone onset and ended at 6000 ms (5800 ms window). The above time window was chosen to obtain a frequency resolution (0.1724 Hz) that included a spectral estimate at 41.2 Hz. To obtain the covariance matrices, we used 124 electrodes (excluding the 4 ocular sensors). To obtain a robust computation of the RESS signal, all four conditions were concatenated across trials within each participant and used to calculate the spatial filter. This spatial filter was applied to the time series of each single trial using the dot product between the filter weights and the trial’s electrode by time matrix. This resulted in a single time series per condition and participant, with the electrode dimension removed. The output was then transformed into the frequency domain by means of Discrete Fourier Transform (fft.m in Matlab). Spectral power was estimated as the absolute value of the complex Fourier coefficients after windowing each time series with a cosine square function (20 points rise/fall) and normalizing by the number of input time points. Then, the power at the target frequency of 41.2 Hz (ASSR) was extracted, using the same time window of analyses (5800 ms) as for the RESS computation, and the signal-to-noise ratio (SNR) spectrum was computed according to the the RESS example script in Cohen and Gulbinaite (2017). The spatial filter was also retained to later identify which electrodes contributed the most to the filter weights.

### Alpha-band power

A family of Morlet wavelets were convolved with the artifact-free EEG trials to obtain the power information across time for the brain oscillations between 3.03 and 55 Hz, in frequency steps of 0.15. The Morlet coefficient was set at 10 allowing a suitable trade-off between frequency smearing and temporal smearing at the target frequency of 10 Hz (Keil et al., 2022): The frequency uncertainty at 10 Hz (center of the alpha band) was 1 Hz and the temporal uncertainty was 1.571 seconds. After obtaining the time-frequency information for each trial, the single-trial data were averaged separately across the trials of each condition. A baseline adjustment was applied by dividing the entire trial segment (-600 to 6000 ms) for each electrode and frequency by the average of the pre-stimulus segment between -500 and -100 ms, and expressing the resulting ratios as percent change relative to baseline. Finally, for the statistical analyses, alpha-band power was obtained by averaging the frequencies between 9.39 to 11.51 Hz.

### Pupillometry

Pupil diameter was measured using an EyeLink 1000 Plus eye tracker system equipped with a 16 mm lens. The data were sampled at a rate of 500 Hz and the pupil diameter was calculated by fitting an elliptic shape to the pupil’s mass threshold. To calibrate and validate the eye tracker, participants were asked to follow a white circle with their gaze on a twelve-point grid. Data that were lost due to eyeblinks were interpolated during the pre-processing stage. The pupil data were then filtered using a 7.5-1500 Hz Butterworth filter and processed from 600 milliseconds before to 6000 milliseconds after the tone onset. In the last stage of pre-processing, the data were converted from arbitrary units to millimeters. 11 participants with high-misophonia and 6 with low-misophonia symptoms were exclude from the analyses because they had less than 50% of good trials. Finally, the last second of the experimental trial was chosen to average the pupil data and use it for the statistical analyses.

### Statistical analyses

For the current study, a Bayesian bootstrap was implemented, following the methodological approach in Ahumada et al. (2025), for the dot product between the experimental data in the four studied variables (ratings, ASSR amplitude, alpha-band power, and pupillometry) and predetermined model weights (discussed below), which are displayed in Figure 2. These analyses are based on bootstrapped Bayes Factors (Efron, 2011; Rubin, 1981), where posterior distributions of the dot products between the data and the models, and between the data and a null model (scrambled weights) are obtained through bootstrapping the original sample. Dot products are a simple approach to quantify the correspondence between a given hypothesis and empirical data. Specifically, the dot product between data and model weights represents how similar a theoretical model, based on a specific hypothesis, is to the empirical data. Greater and positive dot products indicate greater similarity.

**Figure 2.**
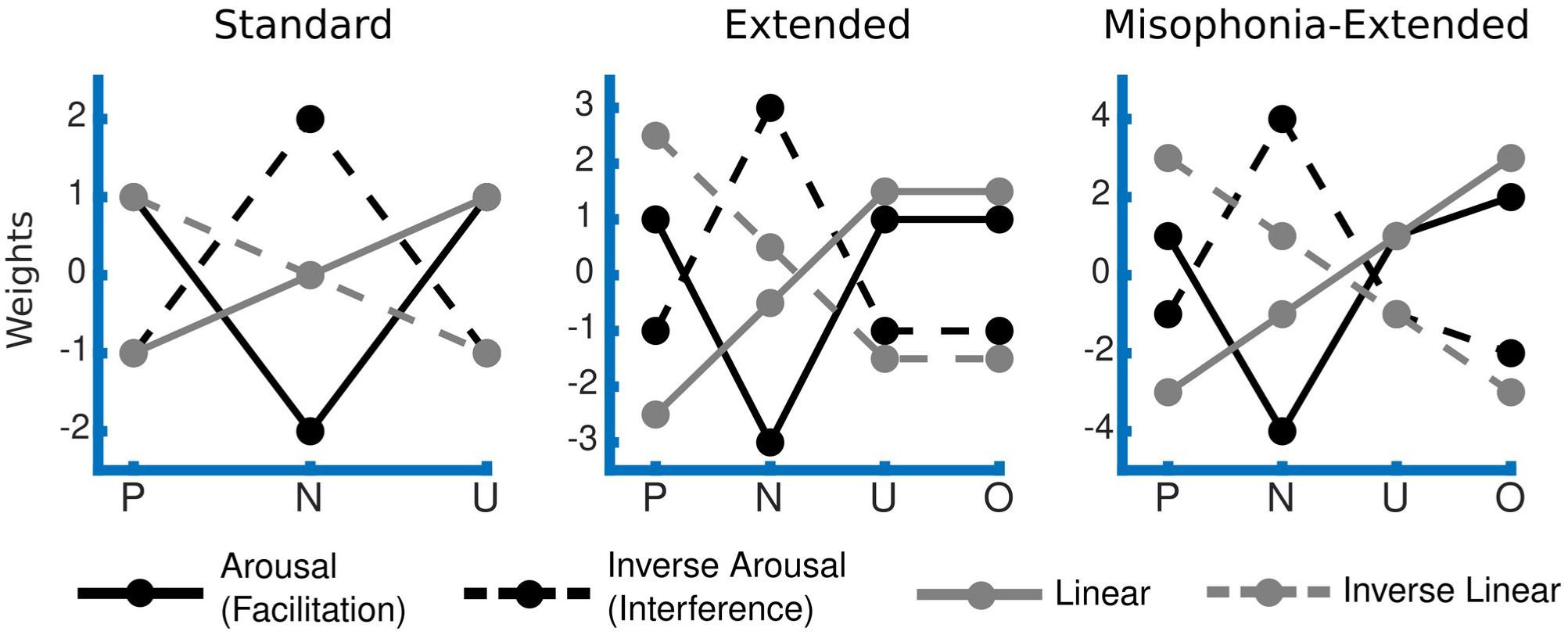
Theoretical Models. *Note.* The weights of these theoretical models were used to compute the dot products together with the empirical data in the four experimental variables (ratings, ASSR, alpha-band power, and pupil diameter) to obtained bootstrapped distributions similarity. The x-axis labels correspond to the first letter of the sound categories’ name: P = Pleasant, N = Neutral, U = Unpleasant, O = Orofacial.

Distributions for the Bayesian analyses were generated through bootstrapping: For each iteration of the bootstrap, participant data for the experimental conditions were randomly selected with replacement and averaged. The resulting grand mean of each condition was then multiplied by the transposed model weights and the null model’s scrambled weights, separately, to obtain the dot product of each one. This iteration is repeated 5000 times.

To examine whether self-reported arousal ratings for the standard naturalistic sound categories (pleasant, neutral, unpleasant) followed an arousal trend, where emotional stimuli are reported as more arousing, a standard arousal model (weights = [1 -2 1], for pleasant, neutral, unpleasant sounds, respectively) was used to compute the dot product in each group (LMS and HMS). In the cases of the valence ratings, a standard linear model (weights = [-1 0 1]) was utilized to evaluate whether the scores of the standard naturalistic sounds displayed a linear increment in rated displeasure. Additionally, to assess the models similarity with the scores of the fourth categories of naturalistic sounds (including the orofacial), an extended arousal (weights = [1 -3 1 1]), misophonia-extended arousal (weights = [1 - 4 1 2]), extended linear (weights = [-2.5 -.5 1.5 1.5]), and misophonia-extended linear (weights = [-3 -1 1 3]) models were also evaluated. The extended versions of these models are based on the hypotheses that low misophonia participants will assess the orofacial sounds equally as they assess the unpleasant sounds, whereas individuals endorsing misophonia symptoms will evaluate the orofacial sounds as more arousing and unpleasant than the rest of the sound categories.

To test whether the naturalistic sounds affected the tone’s ASSR amplitude, the dot product between the amplitude and competition-based models were calculated. As mentioned previously, one possible outcome of competition is represented in the interference model. In this case, we would anticipate a biased processing toward the most significant stimulus, where the ASSR amplitude is reduced in the presence of emotional sounds. The second possible outcome is represented in the facilitation model. In this model, concurrent stimuli are amplified, boosting the ASSR amplitude when the tone is accompanied by emotional sounds. To assess these theories, three weights of the interference and facilitation models were utilized to test not only the effect of competition in standard emotional categories (pleasant, neutral, and unpleasant), but also the effect of competition with symptom specific stimuli (orofacial sounds). Similar to the rating analysis, the standard interference model uses the weights [-1 2 -1], the extended interference model uses [-1 3 -1 -1], and the misophonia-extended model employs [-1 4 -1 -2]. The three facilitation models have the opposite weights of the interference models (see Figure 2); consequently, whenever one model fits well the data, the other model statistic will not be reported.

To analyze the pupil and alpha-power data, bootstrapped distributions of dot products were calculated between the empirical data and each of the models—including three versions of the arousal model (standard, extended, and misophonia-extended) and three versions of the linear model (standard, extended, and misophonia-extended). Inverse versions of these models were also tested (standard inverse arousal, extended inverse arousal, inverse misophonia-extended arousal, standard inverse linear, extended inverse linear, misophonia-extended inverse linear) to determine whether effects beyond arousal and displeasure influenced these variables. As with the facilitation and interference models tested on the ASSR data, only the best-fitting model will be reported, as it is assumed that the opposing version underperformed.

In the case of the alpha-band power statistical analyses, the ∼10 Hz frequency was averaged across time in time bins of one second each after the naturalistic sound onset. The last second was not added to the analyses to avoid the edge artifact and the motor preparation response to the ratings (for trials in the second block). The above resulted in four time-bins, in which the alpha-band power was averaged (Bin 1 = 1:1000 ms, Bin 2 = 1001:2000 ms, Bin 3 = 2001:3000 ms, and Bin 4 = 3001:4000 ms). Taking into account the general activity of alpha across all electrodes and the entire time window, three electrode clusters were selected to further average alpha-band and then apply the Bayesian bootstrapping. In the 128-channel Geodesic net, the left-central cluster corresponded to the electrodes 29, 30, 35, 36, 37, and 41; right-central cluster to 87, 103, 104, 105, 110, and 111; and parietal cluster to 62, 67, 71, 72, 76, and 77. Therefore, alpha-band power was obtained for each of the naturalistic conditions and groups, in four time bins after sound onset and in three electrode clusters. Then, the previously mentioned models were used on the segmented and clustered alpha-band data to produce the bootstrapped distributions.

Finally, after bootstrapping the average dot products for each variable and within each experimental group, Bayes Factors were calculated with the resulted posterior distribution of dot products. Specifically, posterior odds are calculated using the likelihood of the dot product between the variables of interest and the theoretical models; while the prior odds are calculated using the likelihood of the dot product between the variables of interest and the null model. For both cases, the priors were set to 1 (uninformative priors) and BFs were calculated as the ratio of posterior over prior odds under the assumptions that the dot products from the data against the models are greater than the dot products from the data against the null model (BF_10_ ^fit^). Additionally, to estimate group differences, Bayes Factors were calculated between the bootstrapped similarity distributions for the LMS and HMS groups. The odds ratio between the LMS distribution over the HMS distribution is presented as BF_10_ ^groups^, while the opposite is presented as BF_01_^groups^. In the case of the analyses in the alpha-band frequency, average transitive Bayes Factors were obtained across the time bins and the best-fitting models were reported for each electrode cluster and group. Transitive Bayes Factors are obtained by dividing a model’s BF_10_ ^fit^ (with values above 3) by another model’s BF_10_^fit^. For instance, for those models that quantify the trend of the first three naturalistic sounds (pleasant, neutral, and unpleasant), the average BF (across the time bins) of the standard arousal/inverse arousal was divided by the BF of the standard linear/inverse linear model. The Bayes Factors were interpreted according to Jeffreys’ guidelines for evidence strength (Jeffreys, 1998).

## Results

### Hyperacusis

In a first step, we examined the extent to which test LDLs, used as a proxy of hyperacusis, differed between the low-misophonia (LMS, N = 31) and high-misophonia (HMS, N = 29) groups, independent-samples t-tests were performed across LDL thresholds of five tones with different pitches (see Methods), and the sum of the thresholds from all tones. Results showed no significant differences regarding the loudness thresholds between the groups, in the first tone (LMS [M = 5.48, SD = 3.60], HMS [M = 6.41, SD = 3.85], t_(56.98)_ = -0.96, p = 0.34); the second tone (LMS [M = 4.77, SD = 3.38], HMS [M = 5.65, SD = 4.04], t_(54.80)_ = -0.91, p = 0.37); the third tone (LMS [M = 4.19, SD = 3.26], HMS [M = 5.48, SD = 4.00], t_(54.13)_ = -1.36, p = 0.18); the fourth tone (LMS [M = 4.52, SD = 3.66], HMS [M = 5.31, SD = 3.80], t_(57.48)_ = -0.82, p = 0.42); the fifth tone (LMS [M = 3.29, SD = 3.02], HMS [M = 4.41, SD = 3.48], t_(55.63)_ = -1.33, p = 0.19); or the total threshold (LMS [M = 22.26, SD = 16.05], HMS [M = 27.26, SD = 17.96], t_(56.19)_ = -1.14, p = 0.26).

### Ratings

Figure 3A shows the averaged self-reported arousal and valence (displeasure) for both groups and for each of the sound categories. Figure 3B displays the bootstrapped distributions of the dot products between data and model weights. In the case of the arousal ratings, the bootstrapped similarity values showed strong support for the standard arousal model than for the null model, in both the LMS (BF_10_^fit^ = 17,998.00) and the HMS (BF_10_^fit^ = 634.61) groups. No evidence for group differences in arousal ratings was found (BF_10_ ^groups^ = 0.51). When including the fourth sound category (orofacial), there was decisive evidence for the extended arousal model in both the LMS (BF_10_ ^fit^ = 167,650.00) and HMS (BF_10_^fit^ > 1’000,000) groups. There was moderate evidence that the groups were different (BF_01_^groups^ = 8.83), indicating a better fit of the extended arousal model in the HMS group, compared to the LMS group. The last model that was tested against the arousal scores was the misophonia-extended arousal model. There was decisive evidence in favor of this model in the LMS (BF_10_ ^fit^ = 293,140.00) and the HMS (BF_10_ ^fit^ > 1’000,000) group. Again, there was strong evidence of a better fit in the HMS group compared to the LMS group (BF_01_ ^groups^ = 14.37). Transitive Bayes Factors suggest that, in the LMS group, the extended and misophonia-extended models did not differ in their goodness of fit (BF^transitive^ = 1.75). However, in the HMS group, the misophonia-extended arousal model was a better fit than the extended model (BF^transitive^ = 28.80).

**Figure 3.**
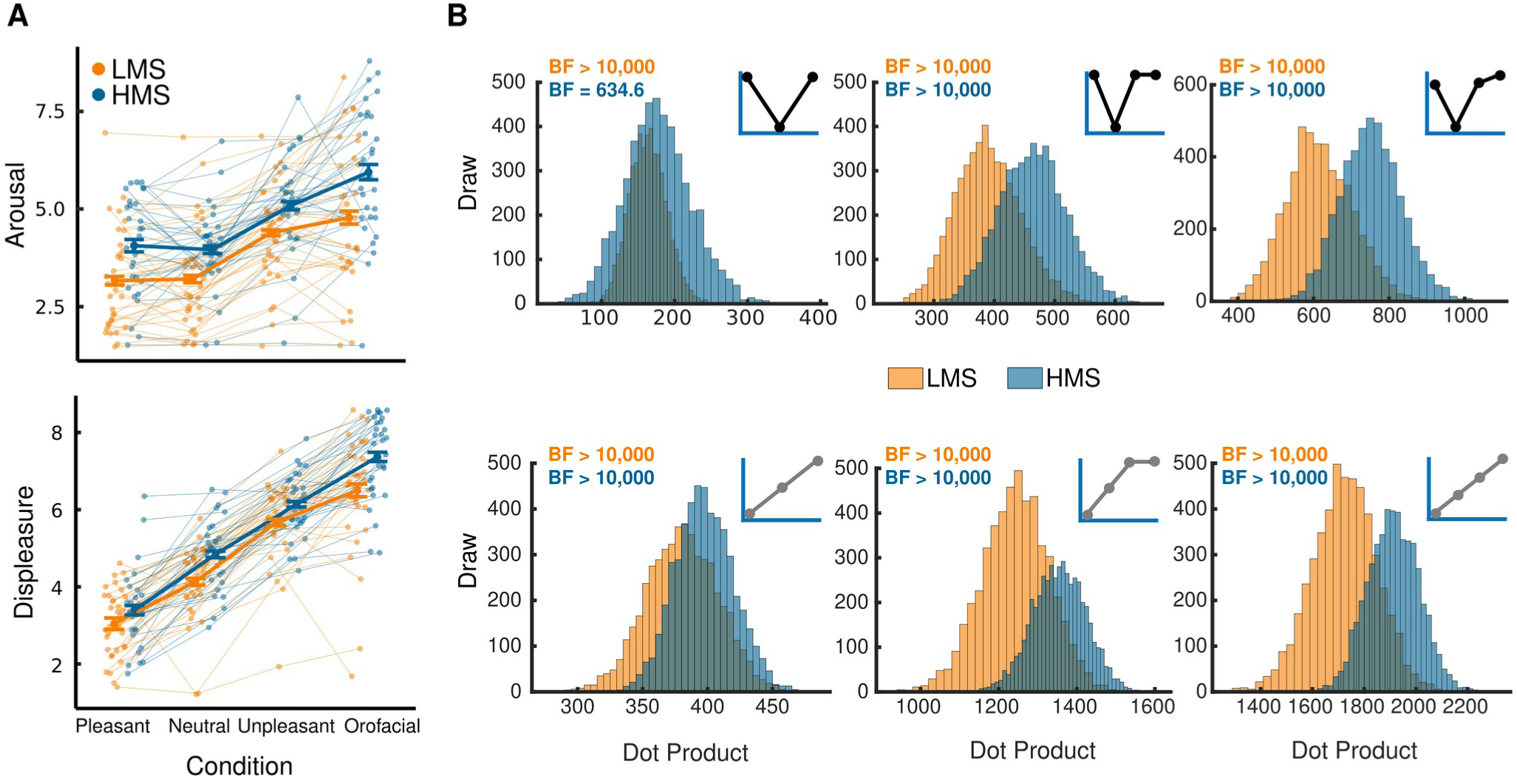
Subjective Ratings of Arousal and Valence (Displeasure) for Naturalistic Sounds. *Note.* **A:** Average arousal and valence (displeasure) ratings for each sound category (pleasant, neutral, unpleasant, and orofacial) in the LMS and HMS groups, including individual ratings. Error bars represent within-subject standard errors. **B (top):** Bootstrapped distributions of dot products between the arousal ratings and the arousal (standard, extended, and misophonia-extended) models in the LMS and HMS groups. **B (bottom):** Bootstrapped distributions of dot products between the valence ratings and the linear (standard, extended, and misophonia-extended) models in the LMS and HMS groups. BF values depict the goodness of fitting of the models for each empirical group (orange = LMS, blue = HMS).

The linear model tested against the valence ratings indicated that this model fit both groups better than a null model (both BF ^fit^ >= 1’000,000). There was moderate evidence supporting group differences in valence ratings (BF_01_^groups^ = 3.01), with the model fitting the HMS group better than the LMS group. When including the fourth sound category, the fit of the extended linear model on the valence ratings was again better than a null model in both groups (both BF_10_^fit^ >= 1’000,000). Once again, there was moderate to strong evidence supporting that groups differed (BF_01_ ^groups^ = 10.86), with greater support for the extended model in the HMS group than in the LMS group. Lastly, the misophonia-extended linear model also fit the valence ratings in the LMS (BF_10_^fit^ > 1’000,000) and HMS (BF_10_^fit^ > 1’000,000) groups, with strong evidence that the model provided a superior fit for the HMS group compared to the LMS group (BF_01_ ^groups^ = 16.50). Comparing the models that included the orofacial conditions, we observed that, in the LMS group, the extended linear model fit the data better than the misophonia-extended version (BF^transitive^ = 32.12). While in the HMS, the misophonia-extended linear model aligned to the data better than the extended version (BF^transitive^ = 7,187.14).

### ASSR

Figure 4A illustrates the isolation of the ASSR using the RESS method. The topography represents the estimated scalp locations that contributed to the spatial filters for the RESS component corresponding to the ASSR. The average of the ASSR amplitude across condition (sound categories) and experimental group is shown in Figure 4B, while Figure 4C shows the bootstrapped distributions of the dot products (the facilitation models against the empirical data).

**Figure 4.**
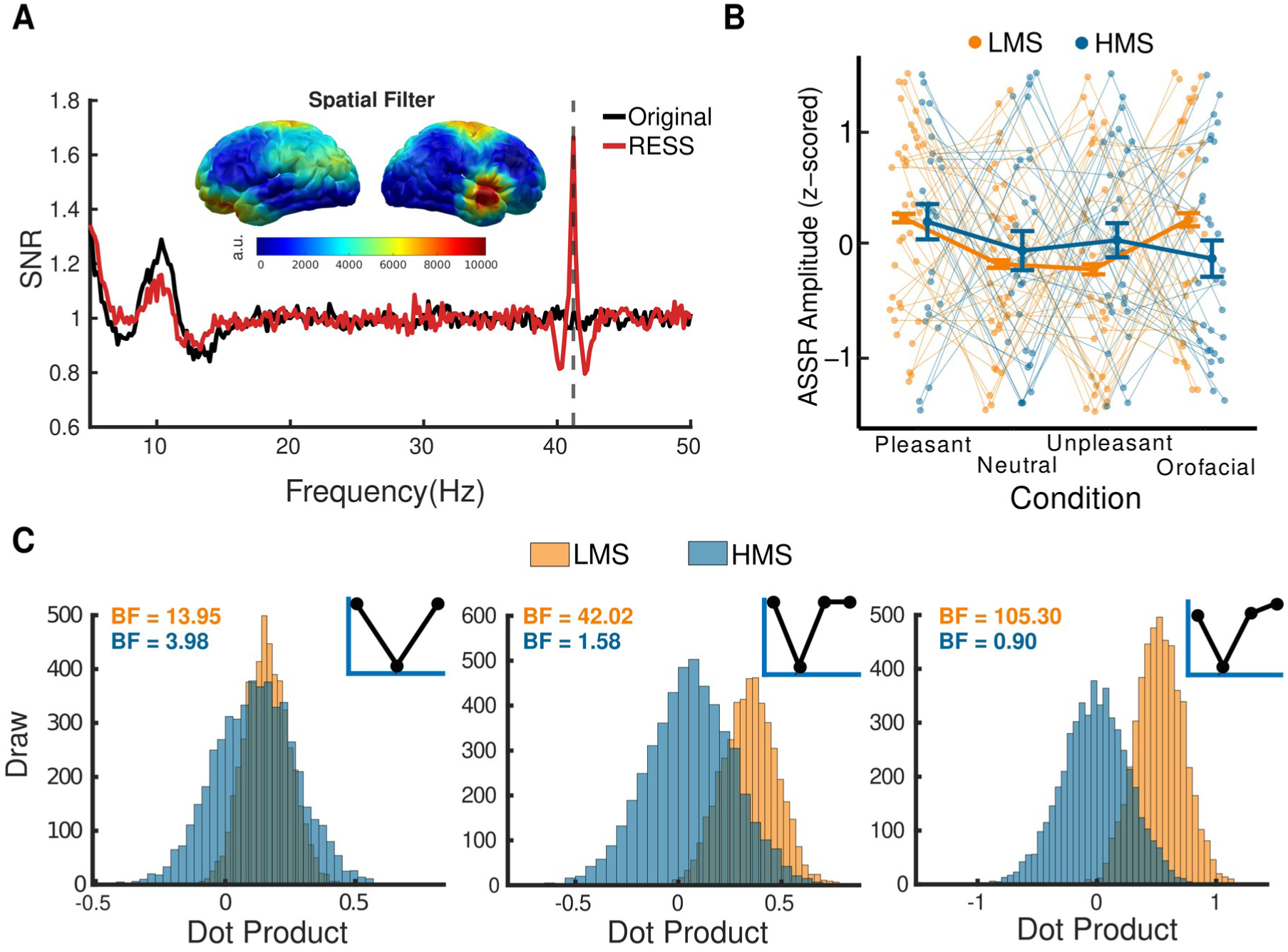
ASSR Amplitude. *Note.* Changes in auditory steady-state response (ASSR) amplitude. **A:** Spectrogram signal-to-noise ratio (SNR) showing the enhancement of the ASSR signal after applying the RESS solution. The topographic map illustrates the estimated electrode locations contributing to the spatial filters. **B:** Average z-transformed ASSR amplitude for each condition and group, including individual data points. Error bars represent within-subject standard errors. **C:** Bootstrapped distributions of dot products for the facilitation models (standard, extended, and misophonia-extended) in the LMS and HMS groups. BF values depict the goodness of fitting of the models for each empirical group (orange = LMS, blue = HMS).

It was found that the facilitation model was a better fit for the ASSR in the LMS group than a null model (BF_10_ ^fit^ = 13.95). In the case of the HMS group, the same model fit the data moderately better than a null model (BF_10_ ^fit^ = 3.98). There was not enough evidence supporting the hypothesis that the groups were different in their ASSR on this model (BF_10_ ^groups^ = 1.88). The extended facilitation model with the orofacial condition showed a better fit than the null model in the LMS group (BF_10_ ^fit^ = 42.02), but not the HMS group (BF_10_^fit^ = 1.58). This group difference was meaningful (BF_10_ ^groups^ = 18.54). Regarding the fitting of the misophonia-extended facilitation model, decisive evidence showed a better fit for this model compared to a null model, for the data from the LMS distribution (BF_10_^fit^ = 105.30). However, this model failed to fit the data from the HMS group (BF_10_ ^fit^ = 0.90). Again, there was strong evidence showing that this group difference was meaningful (BF_10_^groups^ = 49.30). Although the Bayes Factor for the misophonia-extended facilitation model was greater than the Bayes Factor for the extended facilitation model in the LMS group, the difference between these models was weak (BF^transitive^ = 2.51).

Follow-up analyses further highlighted that the primary source of differences between the two groups was the ASSR response during orofacial sounds, with strong evidence in support of the notion that the tone-evoked ASSR amplitude in the LMS group was greater than in the HMS group when embedded in orofacial sounds (BF_10_^groups^ = 151.06).

### Alpha-band power

Figure 5A shows the baseline-adjusted time-varying power during the listening paradigm, averaged across conditions. As mentioned above, four time bins were used to characterize the time course of alpha-power changes in response to the compound stimuli, in three different electrode clusters (left central, right central, and parietal cluster). The grand mean alpha power change values in each time bin, electrode cluster, and group are shown in Figure 5B. For a better visualization, the probability distribution function (PDF) corresponding to each bootstrap distribution for the best-fitting models, across electrode clusters and experimental groups, were plotted in Figure 5C. The PDFs were calculated using the fitdist function in Matlab. Besides, the Bayes Factors for each model fit to the alpha-band changes are reported in Table 2. Whenever an opposite model fit the data better (reaching a BF_10_ above 3), its corresponding Bayes Factor was reported in the table within brackets.

**Figure 5.**
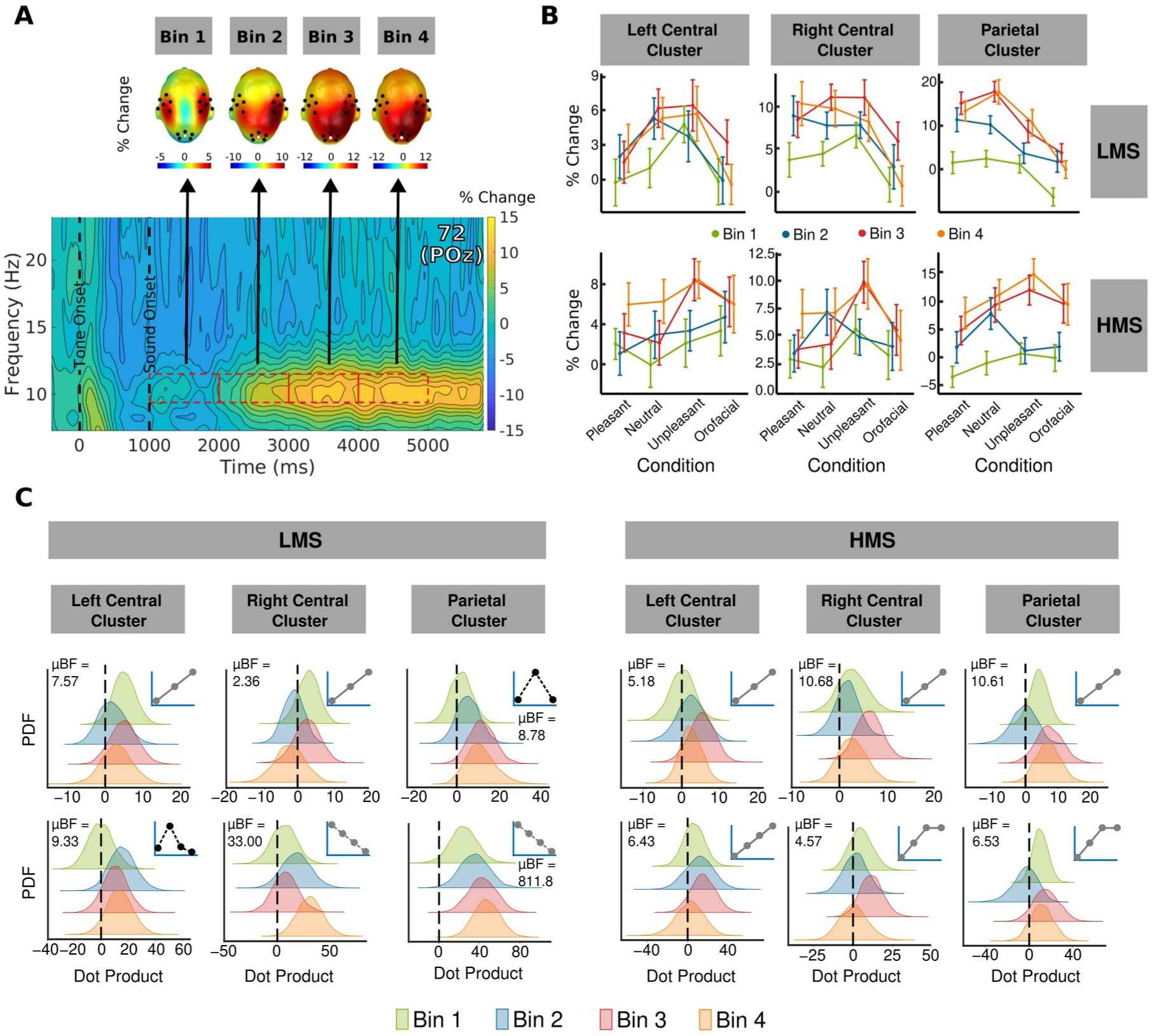
Alpha-band power. *Note.* Alpha-band power fluctuations in response to naturalistic sounds. **A:** Time-frequency spectrogram illustrating the increase in alpha-band power following naturalistic sound onset. Four 1- second time bins (indicated by red-dashed rectangles) were selected for model testing across three electrode clusters (marked by black dots on the topographic maps). **B:** Average percentage change in alpha-band power for each condition and group, calculated across electrode clusters and time bins. Error bars represent within-subject standard errors. **C:** Bootstrapped distributions of dot products for the best-fitting models, transformed into probability density functions (PDFs) for visualization, and shown separately for the LMS and HMS groups across electrode clusters and time bins. Averaged BF values across time-bins depict the goodness of fitting of the models.

**Table 2.**
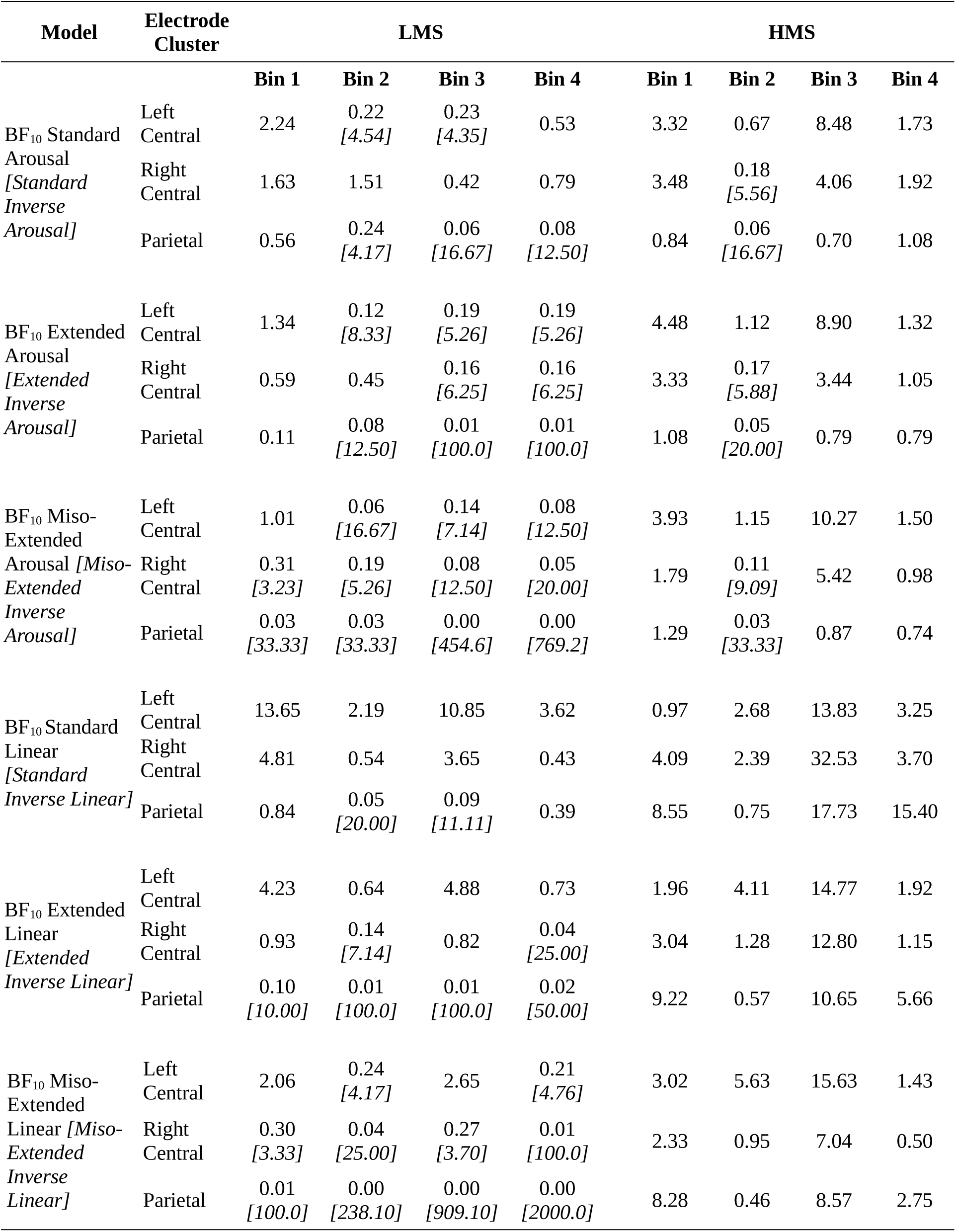
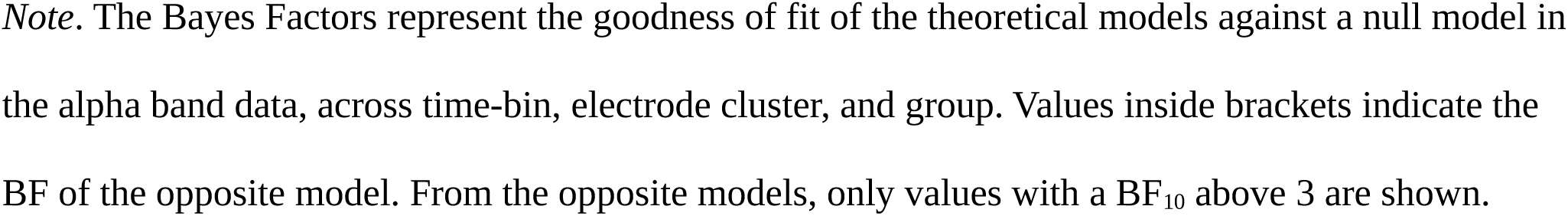
Bayes Factors for Goodness of Fit of Theoretical Models in Alpha-Band Power.

| Model | Electrode Cluster | LMS |  |  |  | HMS |  |  |  |
| --- | --- | --- | --- | --- | --- | --- | --- | --- | --- |
|  |  | Bin 1 | Bin 2 | Bin 3 | Bin 4 | Bin 1 | Bin 2 | Bin 3 | Bin 4 |
| BF<sub>10</sub> Standard Arousal<br>*[Standard Inverse Arousal]* | Left Central | 2.24 | 0.22<br>[4.54] | 0.23<br>[4.35] | 0.53 | 3.32 | 0.67 | 8.48 | 1.73 |
|  | Right Central | 1.63 | 1.51 | 0.42 | 0.79 | 3.48 | 0.18<br>[5.56] | 4.06 | 1.92 |
|  | Parietal | 0.56 | 0.24<br>[4.17] | 0.06<br>[16.67] | 0.08<br>[12.50] | 0.84 | 0.06<br>[16.67] | 0.70 | 1.08 |
| BF<sub>10</sub> Extended Arousal<br>*[Extended Inverse Arousal]* | Left Central | 1.34 | 0.12<br>[8.33] | 0.19<br>[5.26] | 0.19<br>[5.26] | 4.48 | 1.12 | 8.90 | 1.32 |
|  | Right Central | 0.59 | 0.45 | 0.16<br>[6.25] | 0.16<br>[6.25] | 3.33 | 0.17<br>[5.88] | 3.44 | 1.05 |
|  | Parietal | 0.11 | 0.08<br>[12.50] | 0.01<br>[100.0] | 0.01<br>[100.0] | 1.08 | 0.05<br>[20.00] | 0.79 | 0.79 |
| BF<sub>10</sub> Miso-Extended Arousal<br>*[Miso-Extended Inverse Arousal]* | Left Central | 1.01 | 0.06<br>[16.67] | 0.14<br>[7.14] | 0.08<br>[12.50] | 3.93 | 1.15 | 10.27 | 1.50 |
|  | Right Central | 0.31<br>[3.23] | 0.19<br>[5.26] | 0.08<br>[12.50] | 0.05<br>[20.00] | 1.79 | 0.11<br>[9.09] | 5.42 | 0.98 |
|  | Parietal | 0.03<br>[33.33] | 0.03<br>[33.33] | 0.00<br>[454.6] | 0.00<br>[769.2] | 1.29 | 0.03<br>[33.33] | 0.87 | 0.74 |
| BF<sub>10</sub> Standard Linear<br>*[Standard Inverse Linear]* | Left Central | 13.65 | 2.19 | 10.85 | 3.62 | 0.97 | 2.68 | 13.83 | 3.25 |
|  | Right Central | 4.81 | 0.54 | 3.65 | 0.43 | 4.09 | 2.39 | 32.53 | 3.70 |
|  | Parietal | 0.84 | 0.05<br>[20.00] | 0.09<br>[11.11] | 0.39 | 8.55 | 0.75 | 17.73 | 15.40 |
| BF<sub>10</sub> Extended Linear<br>*[Extended Inverse Linear]* | Left Central | 4.23 | 0.64 | 4.88 | 0.73 | 1.96 | 4.11 | 14.77 | 1.92 |
|  | Right Central | 0.93 | 0.14<br>[7.14] | 0.82 | 0.04<br>[25.00] | 3.04 | 1.28 | 12.80 | 1.15 |
|  | Parietal | 0.10<br>[10.00] | 0.01<br>[100.0] | 0.01<br>[100.0] | 0.02<br>[50.00] | 9.22 | 0.57 | 10.65 | 5.66 |
| BF<sub>10</sub> Miso-Extended Linear<br>*[Miso-Extended Inverse Linear]* | Left Central | 2.06 | 0.24<br>[4.17] | 2.65 | 0.21<br>[4.76] | 3.02 | 5.63 | 15.63 | 1.43 |
|  | Right Central | 0.30<br>[3.33] | 0.04<br>[25.00] | 0.27<br>[3.70] | 0.01<br>[100.0] | 2.33 | 0.95 | 7.04 | 0.50 |
|  | Parietal | 0.01<br>[100.0] | 0.00<br>[238.10] | 0.00<br>[909.10] | 0.00<br>[2000.0] | 8.28 | 0.46 | 8.57 | 2.75 |

*Note.* The Bayes Factors represent the goodness of fit of the theoretical models against a null model in the alpha band data, across time-bin, electrode cluster, and group. Values inside brackets indicate the BF of the opposite model. From the opposite models, only values with a $BF_{10}$ above 3 are shown.

Overall, the analysis of alpha-band activity in the LMS group revealed distinct trends across different clusters. When considering only the standard sounds (pleasant, neutral, and unpleasant), the left central cluster showed a strong fit with the standard linear trend (µBF10 = 7.57). In contrast, the right central cluster did not show a clear fit with any of the tested models, although anecdotal evidence suggested a moderate fit with the standard linear model (µBF10 = 2.36). The parietal cluster, on the other hand, was best described by the standard inverse linear model (µBF10 = 8.78). When including all four sound categories, the misophonia-extended inverse arousal model fit the data well across the time bins in the left central cluster (µBF_10_ = 9.33); while the misophonia-extended inverse linear model fit the data better in the right central (µBF_10_ = 33.00) and parietal bin (µBF_10_ = 811.80).

Regarding the HMS group, the standard linear model aligned better with the alpha activity in the left central (µBF_10_ = 5.18), right central (µBF_10_ = 10.68), and parietal cluster (µBF_10_ = 10.61) when considering only the standard sounds. After taking into account the four sound categories, the misophonia-extended linear model fit the alpha-band trend better in the left central cluster (µBF_10_ = 6.43); while the extended linear model fit the data better in the right central (µBF_10_ = 4.57) and parietal cluster (µBF_10_ = 6.53).

### Pupilometry

Changes in pupil diameter over time and across the naturalistic sound categories and experimental groups can be observed in Figure 6A. Individual pupil data were z-transformed for visualization (Figure 6B). As for alpha-power, we used the arousal and linear models to evaluate whether the data from both groups follow those trends, see Figure 6C. We found that the standard arousal model did not fit the bootstrapped similarity distributions for the LMS group (BF_10_^fit^ = 0.21). Instead, the data moderately fit the inverse of the standard arousal model (BF_10_ ^fit(inverse^ ^arousal)^ = 4.57).

**Figure 6.**
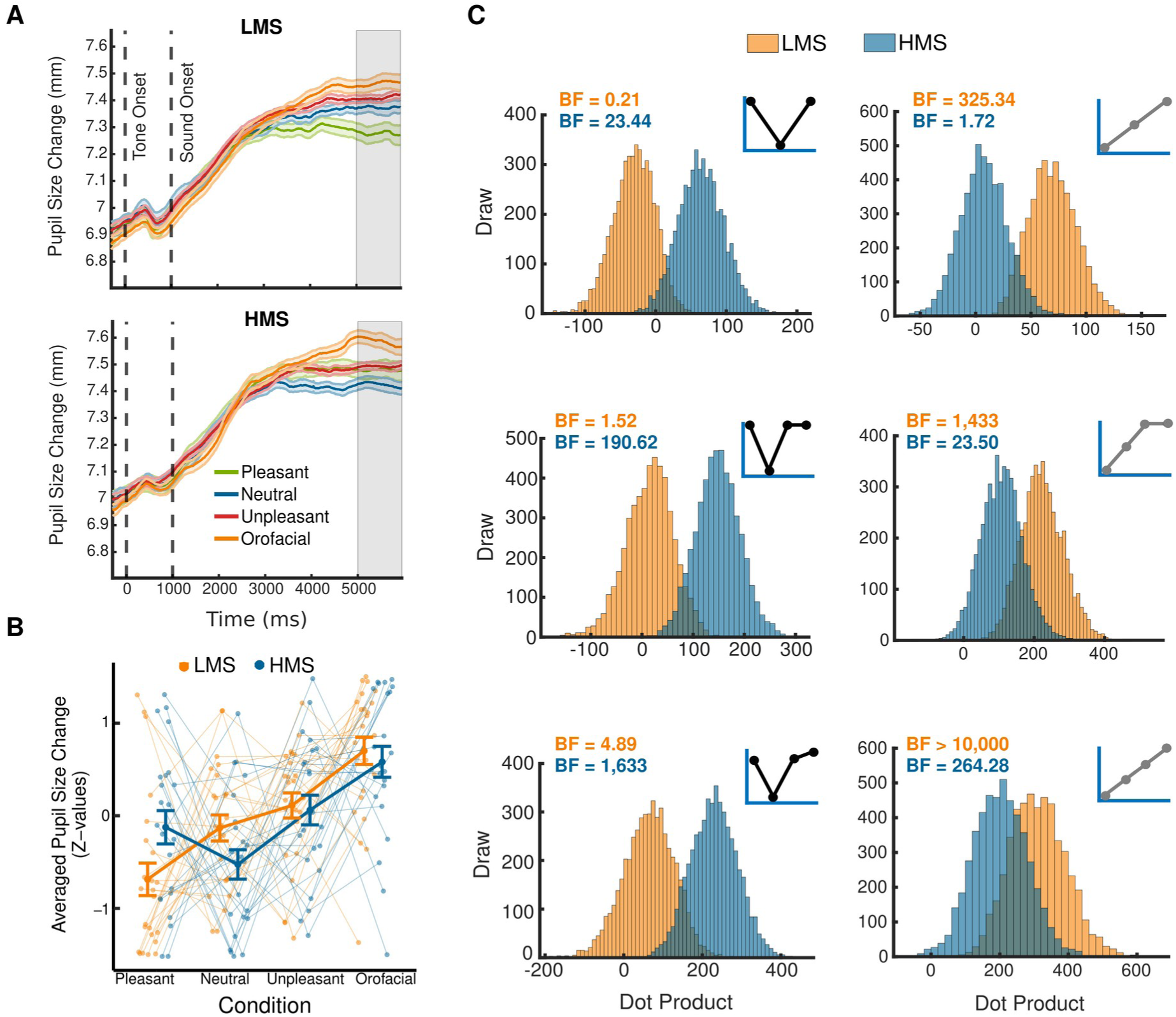
Pupil Dilation. *Note.* Pupil size changes in response to tone and naturalistic sounds. **A:** Time course of pupil size changes following tone and naturalistic sound onset, shown for each condition and group. The gray area highlights the last second of each trial, which was used to calculate the average pupil size for statistical analysis. **B:** Average z-transformed pupil size per condition and group, including individual data points. Error bars represent within-subject standard errors. **C:** Bootstrapped distributions of dot products for arousal (standard, extended, and misophonia-extended) and linear (standard, extended, and misophonia-extended) models in the LMS and HMS groups. BF values depict the goodness of fitting of the models for each empirical group (orange = LMS, blue = HMS).

There was strong evidence that the standard arousal model fit the data from the HMS group better than a null model (BF_10_ ^fit^ =23.44). Consequently, strong evidence supported that the groups were different (BF_01_^groups^ =261.14), with the HMS group showing a stronger resemblance to the standard arousal model. After adding the fourth stimulus category to the model comparison, the extended arousal model failed to fit the data from the LMS group (BF_10_^fit^ = 1.52), but strongly fit the data from the HMS group (BF_10_^fit^ = 190.62) compared to a null model. Strong evidence in favor of a difference between both groups was found again (BF_01_^groups^ = 174.55). The last model tested was the misophonia-extended arousal model. This model had moderate support in the LMS group (BF_10_^fit^ = 4.90), and substantial support in the HMS group (BF_10_^fit^ = 1,633.10). Again, there was decisive evidence showing that pupil dilation in the HMS group was more similar to the misophonia-extended arousal model than the LMS groups. The HMS group’s pupil dilation patterns were more consistent with the misophonia-extended arousal model than the LMS groups, with decisive support (BF_01_^groups^ = 112.41).

The linear/inverse linear models were also tested on the pupil data. The standard linear model fit well the data from the LMS group (BF_10_^fit^ = 325.34), but not the data from the HMS goup (BF_10_^fit^ = 1.72). There was decisive evidence that these group differences were meaningful (BF_10_ ^groups^ = 156.25). The extended linear model fit the LMS (BF_10_ ^fit^ = 1,432.80) and the HMS (BF_10_ ^fit^ = 23.50) groups, with strong support in favor of the group differences being meaningful, with better model alignment with the LMS data than the HMS data (BF_10_ ^groups^ = 24.75). Finally, the misophonia-extended linear model fit the data in the LMS (BF_10_ ^fit^ = 17,279.00) and HMS (BF_10_ ^fit^ = 264.28) groups. Moderate evidence revealed that pupil dilation in the LMS group was more closely aligned with this model than the HMS group (BF_10_^groups^ = 9.35). Notably, the models that fit the pupil data best in the LMS group were the three versions of the linear model. In particular, the misophonia-extended linear model fit the pupil data better than the extended model, when the four sound categories were tested (BF^transitive^ = 12.06). In the case of HMS group, the models that aligned best with the pupil data were the three version of the arousal model, with the misophonia-extended model fitting best when considering the four sound categories (BF^transitive^ = 8.57).

The previously mentioned Bayes Factors for the affective ratings, ASSR, and pupil dilation variables can be also observed in the supplementary Table S2 to facilitate the visual inspection on the statistical results. Only the Bayes Factors for the models that were tested on each variable are displayed.

## Discussion

The brain’s ability to select specific subsets of information from multiple auditory streams is an adaptive mechanism for processing information under limited capacity. The present study aimed to isolate the auditory neural response (ASSR) of an amplitude modulated tone that was presented concurrently with naturalistic sounds varying in emotional content. Additional behavioral and physiological variables were measured to assess affective responses to the compound auditory stimuli. This was conducted in two groups of listeners who differed in their endorsing of symptoms of misophonia. A-priori theoretical models were tested for all dependent variables, using Bayesian model fitting procedures. Models were rooted in either interference or facilitation-based theories of how concurrent auditory streams interact. Interference-based views are largely based on studies in the human visual system (De Echegaray et al., 2024; Muller et al., 2008) and on research with intermodal tasks (e.g., Saupe et al., 2009). In contrast, support for facilitation views is based on electrophysiological and behavioral studies of object-based auditory attention and stream integration (Alain & Arnott, 2000; Shinn-Cunningham, 2008). We examined these two broad hypotheses in the context of two specific goals: (1) To characterize the nature of the interaction between an amplitude modulated tone and a concurrent naturalistic sound varying in motivational relevance; and (2) to examine the role of inter-individual differences, namely misophonia symptoms, in these effects.

As expected, ratings of the naturalistic sounds on scales of emotional arousal and hedonic valence varied systematically as a function of sound category across both groups. In addition, individuals endorsing high levels of misophonia symptoms reported more extreme displeasure and arousal to the orofacial sounds than those in the LMS group. This was also true for pupil dilation, where all participants showed overall heightened responses to unpleasant sounds, but HMS individuals showed more extreme sympathetic responses (more dilation) in response to orofacial sounds, supporting the extended misophonia model. Interestingly, the arousal ratings for all sound categories were slightly more arousing in the HMS group than the LMS group. Regarding the pupil dilation, the LMS group showed a linear increase in pupil diameter across sound categories that mirrored the displeasure ratings. The HMS group showed arousal-related condition differences, with even more extreme pupil dilation for the orofacial category. This replicates a recent finding that misophonia trigger sounds prompted heightened pupil diameter increase in misophonia sufferers, but also in control participants (Oszczapinska et al., 2025). Overall, pupil dilation data largely converged with arousal ratings, suggesting that this affective self-reported variable encompass the intensity of activation of the motivational system (Codispoti & De Cesarei, 2007; Keil et al., 2001; Lang et al., 1998).

Changes in alpha EEG power also differed across the sound categories and between the experimental groups. Early in the trial, increased alpha power was seen in centro-lateral electrodes, a projection target of large numbers of auditory cortical neurons (Lu et al., 2022; Picton et al., 2003). Later in the trial, alpha activation expanded to a more parietal location, as previously seen in auditory work with conditioned aversive cues (Farkas et al., 2024; Strauß et al., 2014). Alpha power changes in parietal regions have been attributed to heightened inhibition of irrelevant environmental sound during speech processing (see also Obleser & Weisz, 2012). We provide additional discussion of alpha-power changes in the present study below, in the context of the tone-evoked ASSR.

### Competition evaluation

Differences in the tone-evoked ASSR amplitude as a function of sound category were also observed. In general, there was an increase in the ASSR amplitude in both groups (LMS and HMS) when tones were accompanied by pleasant or unpleasant sounds relative to neutral sounds. These findings support facilitation models of auditory selection. Differences in ASSR amplitude between the control and misophonia group were only found for the orofacial sound condition: When listening to tones embedded in orofacial sounds, HMS participants showed a pronounced ASSR amplitude reduction, consistent with a interference effect specifically associated with this symptom-relevant category.

In the visual domain, studies on sensory competition and its emotional modulation have converged to show that emotionally engaging pictures strongly interfere with the processing of concurrent visual cues (Müller et al., 2008; Wieser & Keil, 2011), and certain auditory cues (Saupe et al., 2009). Furthermore, participants with fearful dispositions towards a stimulus, such as snake phobia (Deweese et al., 2014) or heightened pain sensitivity (Goldway et al., 2022), display stronger interference when viewing fear-relevant stimuli alongside a concurrent stimulus. In the auditory modality, a recent study showed that high-salient but non-emotional soundscapes decreased the electrocortical response (phase-locked activity and gamma power) to concurrent tones, when compared to conditions with mid-salient and low-salient soundscapes (Huang & Elhilali, 2020). As noted above, this type of interference has been interpreted as resulting from a selection process, serving to resolve competition under limited capacity (Desimone & Duncan, 1995).

Competition depends on how limited capacity is managed across various attentional and motivational demands (Franconeri et al., 2013). This includes the extent to which competing stimuli tap into the same sensory system: If two stimuli are processed by two different sensory systems, then the effect of competition often will be minimal (Talsma et al., 2006), but if they are processed by the same sensory system with substantial spatial and temporal overlap, interference may be more likely. In cross-modal or multimodal tasks, the behavioral response and neurocortical activity is often facilitated under cross-modal stimulation (i.e. with audiovisual stimuli) compared to unimodal auditory or visual stimulation (Chen et al., 2016; Kreifelts et al., 2007). For instance, the bilateral posterior superior temporal gyrus (pSTG) and right thalamus were more activated during an audiovisual task compared to independent visual and auditory tasks, and this effect correlated with the enhanced task performance during the audiovisual setting (Kreifelts et al 2007). Thus, facilitation may be observed as a consequence of integrating multimodal information streams. In support of this notion, it has been found that the neural response to task-relevant tones may be relatively amplified when the tone is simultaneously presented with another tone at a different frequency, in a version of the cocktail-party effect (Xiang et al., 2010). Furthermore, detection of naturalistic targets enhances the auditory N2 component when multiple competing sounds were presented together, compared to when presented in isolation (Lewald & Getzmann, 2015). In all of the above cases, neural activation of a target auditory stream was heightened under competition, and this neural effect was related to improved performance, e.g. in auditory detection tasks (Bronkhorst, 2015).

Another line of research dovetails with these considerations. In the context of object-based auditory attention (Shinn-Cunningham, 2008) the present results could be interpreted as integration of the sound and tone into one attended object. This interpretation is also consistent with the fact the feature space of auditory cortex is defined in the frequency domain (Patel & Balaban, 2004), and the tone and sound frequencies overlapped in the present case. Such integration or grouping of features and stimuli into an auditory object has also been found with the cocktail party effect and has been thought to enhance the intelligibility of speech (Bronkhorst, 2015).

Interestingly, and unexpectedly, when comparing the LMS and HMS groups, strong interference was found in the HMS group only when tones were accompanied by orofacial sounds. An extensive literature has established that sensory systems tend to amplify information that is relevant to a participant’s fear, anxiety, depression, and related mental health problems (Wieser & Keil, 2020). This is often referred to as emotion-congruent attentional bias (Bar-Haim et al., 2007; Dennis-Tiwary et al., 2019; Yiend, 2010). Biases have been suggested to arise from modulatory feedback signals from limbic brain structures into sensory areas (Pessoa & Adolphs, 2010; Sabatinelli et al., 2005), or to arise from plastic changes in sensory systems themselves (Li & Keil, 2023). With regards to misophonia, recent research has likewise emphasized a role emotion-modulated physiology in its etiology and maintenance (Kumar et al., 2017), finding for example that the amygdala and the autonomic system, but not the auditory cortex, play a primary role in the manifestation of misophonia (Jastreboff & Jastreboff, 2023). This viewpoint is helpful for explaining heightened pupil dilation and tone-evoked ASSR in response to aversive sounds. It is however notable that in the present study, biased prioritization appeared to result in a relative amplitude <u>decrease</u> for orofacial sounds, specifically in the HMS group.

Alpha power changes were analyzed as an additional electrophysiological measure to shed additional light on findings with the tone-evoked ASSR. Consistent with the idea that lower alpha power indicates active attentive processing of auditory streams (Wisniewski & Zakrzewski, 2023; Yokosawa et al., 2020), the LMS group showed alpha-power modulations that displayed the opposite pattern as the tone-evoked ASSR, especially in the central clusters: Heightened tone-evoked ASSR was accompanied by lower alpha power, especially late in the trial. Parietal alpha power showed a strong decrease for the orofacial sounds, as opposed to pupil diameter responses. By contrast, HMS participants showed a different pattern of alpha modulations, in which specifically unpleasant and orofacial sounds prompted the highest levels of alpha power across time windows and sensor locations. This was consistent with greater focusing on the tone feature as well as with less attention to the aversive and orofacial sounds, potentially reflecting active ignoring processes.

## Conclusions and future directions

The present research did not include a task and relied solely on passive listening. Past work studying competition in the auditory system has found that competition effects tend to be more pronounced when one stimulus is associated with a behavioral response such as a detection task (e.g., Riels et al., 2020). Thus, the present study cannot provide evidence regarding concepts such as active ignoring or emotion regulation. Future work will test these post-hoc hypotheses by including a task and by systematically manipulating processes related to attending and ignoring compound stimuli varying in emotional content.

In summary, the present study provided evidence that the cortical processing of concurrent auditory stimulus streams varying in emotional content is better described by facilitation models than by interference models. Evidence converged to demonstrate heightened auditory cortical processing of tones that were accompanied by emotionally engaging naturalistic sound. Notably, listeners high in misophonia symptoms deviated from this pattern when listening to symptom-relevant orofacial sounds, displaying suppression of the tone-evoked auditory response. In the context of the auxiliary variables that were collected (pupil dilation, affective ratings, alpha power changes), one possible interpretation is that misophonia prompts heightened emotional responses to aversive and especially orofacial sounds. Uniquely to participants with misophonia, this heightened emotional response may not translate into heightened sensory processing and instead may prompt active ignoring or may translate into dysfunctional interference between the tone stimulus and the accompanying naturalistic sound. Future research may disentangle these processes by adding additional controls and behavioral tasks.

