## Supplementary Material for "Resolving Competition in Auditory Cortex: Effects of Emotional Content and Misophonia Sensitivity"

**Table S1**

*Naturalistic Sounds*

| **Pleasant** | **Neutral** | **Unpleasant** | **Orofacial** |
|---|---|---|---|
| 110 (Baby) | 113 (Bows) | 106 (Growl) | Apple chewing |
| 220 (Boy Laughing) | 171 (Country Night) | 281 (Attack) | Peach slurp |
| 351 (Applause) | 322 (Typewriter) | 502 (Engine Failure) | Chewing gum |
| 353 (Baseball) | 376 (Lawnmower) | 624 (Air Raid) | Straw slurp and sigh |
| 809 (Harp) | 925 (Train) | 713 (Sirens) | Crunchy chewing |
| 812 (Choir) | 701 (Fan) | 715 (Alarm) | Speak and lip smack |
| 226 (Laughing) | 132 (Chickens) | 719 (Dentist Drill) | Chewing ice cubes |
| 352 (Sports Crowd) | 374 (Sink) | 732 (Crash) | Speak and chew |
| 817 (Bongos) | 114 (Cattle) | 910 (Electricity) | Breath and drink |

*Note.* All naturalistic sounds, except for the orofacial sounds, were taken from the International Affective Digitized Sounds (IADS-2; Bradley & Lang, 2007).

**Table S2**

*Models' Fitting on the Ratings, ASSR, and Pupil Data*

| Models | Weights | Ratings | | | | ASSR | | Pupil | |
|---|---|---|---|---|---|---|---|---|---|
| | | Arousal | | Valence | | | | | |
| | | **LMS** | **HMS** | **LMS** | **HMS** | **LMS** | **HMS** | **LMS** | **HMS** |
| $BF_{10}$ Standard Arousal (Standard Facilitation) | [-1, 2, -1] | 17,998.00 | 634.61 | NA | NA | 13.95 | 3.98 | 0.21 | 23.44 |
| $BF_{10}$ Extended Arousal (Extended Facilitation) | [-1, 3, -1, -1] | 167,650.00 | > 1M | NA | NA | 42.02 | 1.58 | 1.52 | 190.62 |
| $BF_{10}$ Misophonia-Extended Arousal (Misophonia-Extended Facilitation) | [-1, 4, -1, -2] | 293,140.00 | > 1M | NA | NA | 105.30 | 0.90 | 4.90 | 1,633.10 |
| $BF_{10}$ Standard Linear | [-1, 0, 1] | NA | NA | > 1M | > 1M | NA | NA | 325.34 | 1.72 |
| $BF_{10}$ Extended Linear | [-2.5, -0.5, 1.5, 1.5] | NA | NA | > 1M | > 1M | NA | NA | 1,432.80 | 23.50 |
| $BF_{10}$ Misophonia-Extended Linear | [-3, -1, 1, 3] | NA | NA | > 1M | > 1M | NA | NA | 17,279.00 | 264.28 |
